# Pan-tissue mitochondrial phenotyping reveals lower OXPHOS expression and function across tumor types

**DOI:** 10.1101/2023.06.04.542600

**Authors:** Ilya N Boykov, McLane M Montgomery, James T Hagen, Raphael T Aruleba, Kelsey L McLaughlin, Hannah S Coalson, Margaret A Nelson, Andrea S Pereyra, Jessica M. Ellis, Tonya N Zeczycki, Nasreen A Vohra, Su-Fern Tan, Myles C. Cabot, Kelsey H. Fisher-Wellman

**Author notes:** To whom correspondence should be addressed: Kelsey H. Fisher-Wellman, East Carolina Diabetes and Obesity Institute, 115 Heart Drive, Greenville, NC 27834 USA. Authors contributed equally (Ilya N Boykov, McLane M Montgomery).

## Abstract

Targeting mitochondrial oxidative phosphorylation (OXPHOS) to combat cancer is increasingly being investigated using a variety of small molecule inhibitors. Clinical success for these inhibitors has been hampered due to serious side-effects potentially arising from the inability to discriminate between non-cancerous and cancerous mitochondria. Although mitochondrial oxidative metabolism is essential for malignant growth, mitochondria OXPHOS is also essential to the physiology of all organs, including high-energy-demand organs like the heart. In comparing tumor OXPHOS reliance to these preeminent oxidative organs it is unclear if a therapeutic window for targeting mitochondrial OXPHOS in cancer exists. To address this gap in knowledge, mitochondrial OXPHOS was comprehensively evaluated across various murine tumors and compared to both matched normal tissues and other organs. When compared to both matched normal tissues, as well as high OXPHOS reliant organs like heart, intrinsic expression of the OXPHOS complexes, as well as OXPHOS flux were consistently lower across distinct tumor types. Operating on the assumption that intrinsic OXPHOS expression/function predicts OXPHOS reliance in vivo, these data suggest that pharmacologic blockade of mitochondrial OXPHOS likely compromises bioenergetic homeostasis in healthy oxidative organs prior to impacting tumor mitochondrial flux in a clinically meaningful way. Although these data caution against the use of indiscriminate mitochondrial inhibitors for cancer treatment, considerable heterogeneity was observed across tumor types with respect to both mitochondrial proteome composition and substrate-specific flux, highlighting the possibility for targeting discrete mitochondrial proteins or pathways unique to a given tumor type.

## INTRODUCTION

The possibility that altered mitochondrial function may be integral to cancer biology was first proposed by Otto Warburg^1^. Warburg viewed cancer as an energy problem, hypothesizing that the origins of cancer stem from the inability to generate sufficient ATP energy from respiration^1^. Contrary to Warburg’s ‘energy-centric’ hypothesis, it is now well understood that the role for the mitochondria in cancer extends well beyond its canonical role as an ATP producer^2–4^. For example, metabolite and ROS signals emanating from mitochondria influence critical aspects of cancer physiology, including proliferation, survival, and cell differentiation^5–8^. Moreover, much of the biomass needed for rapid cell division derives from metabolic intermediates produced during mitochondrial oxidative metabolism^9^. Under conditions where mitochondrial oxidative metabolism is severely impaired, such as following genetic deletion of core respiratory complex subunits or pharmacologic inhibition, global disruptions in mitochondrial respiration consistently constrain tumor growth^10–17^. Thus, with a few exceptions (e.g., succinate dehydrogenase mutant tumors^18^), the process of tumorigenesis in mammals appears generally dependent on respiratory competent mitochondria. The apparent requirement for mitochondrial oxidative metabolism to support tumor biology has sparked interest in exploring mitochondrial inhibitors as a therapeutic strategy to limit/revert tumor growth ^19^. Despite pre-clinical efficacy for various mitochondrial inhibitors^15–17,20–22^, progression of experimental mitochondrial inhibitors through clinical trials in cancer has been hampered by dose-limiting toxicity^20,23^, forcing a need to reevaluate the paradigm of targeting mitochondrial oxidative metabolism ^24^.

Although all mitochondria make ATP, the degree to which a given mitochondrial network is organized to accommodate an ATP resynthesis demand varies by tissue type^25,26^. For example, relative to cardiac mitochondria, mitochondria from liver divert fewer resources to protein components of the OXPHOS system^25^. Inter-organ heterogeneity is also evident across the matrix dehydrogenase network that supplies reducing equivalents to support OXPHOS flux^25^. In short, mitochondria across tissues are highly specialized. Such specialization is reflected in both bioenergetic flux, as well as in proteome expression and appears to be dependent on the unique energetic, biosynthetic, and/or signaling demands of a given host cell/tissue/organ^25–30^. Although mitochondrial specialization across mammalian tissues is well described, much less is known regarding how the mitochondrial proteome is organized in cancer. Relative to non-cancer, malignant cells are subjected to unique physiological demands^31^. Based on first principles, accommodating the physiological demands of malignancy should require mitochondrial specialization. Given the critical need for mitochondrial metabolism in cancer, understanding the dynamic interplay between cancer physiology and mitochondrial specialization is essential to the design of mitochondrial-targeted therapeutics that are cancer-specific.

By integrating comprehensive bioenergetic flux analysis with paired analysis of the mitochondrial proteome, herein we present an in-depth characterization of intrinsic OXPHOS functionality across normal mouse tissues and various tumor types. Across tissues (both normal and tumor), percent contribution of the mitochondrial proteome to core energy transduction pathways (matrix dehydrogenase plus OXPHOS) ranged from over 50% in highly oxidative cardiac mitochondria to ∽ 25% in liver tumor mitochondria. Regardless of the primary site of origin, both OXPHOS expression and function were lower in cancer cells compared to normal cells, suggesting a decreased demand for oxidative ATP synthesis in tumors. Moreover, when normalized to a fixed amount of mitochondrial protein (i.e., mitochondrial content), estimated OXPHOS demand (i.e., reliance) across tumor types was observed to be well below that of the body’s preeminent oxidative organs, suggesting that pharmacologic blockade of mitochondrial energy metabolism likely compromises bioenergetic homeostasis in healthy oxidative organs prior to impacting tumor mitochondrial flux in a clinically meaningful way. These observations partly explain findings noted in recent clinical trials; despite pre-clinical success^17^, inhibition of complex I with IACS-010759 recently failed to achieve anti-cancer efficacy in human trials, due primarily to dose-limiting neurotoxicity^23^. Taken together, our results suggest that the therapeutic window for targeting tumor mitochondria using indiscriminate inhibitors of mitochondrial oxidative metabolism is highly constrained by comparably higher OXPHOS reliance in the canonical oxidative organs. Importantly, however, independent of OXPHOS demand/reliance, considerable heterogeneity was observed across tumor types with respect to the stoichiometry of the subunits comprising the OXPHOS complexes, individual dehydrogenase enzymes, as well as SLC25 mitochondrial transporters. These observation highlights the tantalizing opportunity for targeting a given tumor type based on mitochondrial proteins/pathways that are uniquely up or downregulated.

## RESULTS

### Mitochondrial proteome composition does not support an elevated OXPHOS demand in malignant tissues

To begin to understand mitochondrial specialization in cancer, we evaluated mitochondrial proteome composition across mouse tumor types and compared the results to five healthy mouse tissues (brown fat, colon, heart, kidney, and liver). Tumor types selected included tumors from mice exposed to diethylnitrosamine (DEN) to model hepatocellular carcinoma (HCC), as well as azoxymethane/dextran sodium sulfate (AOM/DSS) to model colorectal cancer (CRC). Given that most mitochondrial-targeted drugs in clinical trials target core components of the energy transduction system^20,21,32^, initial analysis was restricted to the OXPHOS complexes (CI, CII, CIII, CIV, CV), as well as the associated dehydrogenase network (DH) (**Fig. 1A**). For each tissue type, data were normalized to total mitochondrial abundance (based on MitoCarta 3.0 database searching^33^), with OXPHOS and DH expression being expressed as a percentage of the total mitochondrial proteome. As noted in a prior publication^25^, the percentage of the mitochondrial proteome dedicated to OXPHOS was highest in the heart (∽34%) and lowest in the liver (∽13%) (**Fig. 1B-C**). Relative to matched normal tissue, percent OXPHOS expression was unchanged in liver tumors and reduced in CRC tumors (**Fig. 1B-C**). Breaking down OXPHOS expression by each complex, heart was once again higher than all other tissues for most complexes, except CII (i.e., succinate dehydrogenase) and CV (i.e., ATP synthase) (**Fig. 1D**). ATP synthase expression was nearly absent in brown adipose (**Fig. 1D**), consistent with brown fat’s role in thermogenesis. In colon tumors, percent CI expression was lower relative to normal colon, suggesting that CI is specifically downregulated in CRC tumors. Percent expression of the OXPHOS complexes was unaltered in liver tumors. Taken together, relative to matched normal, OXPHOS expression was either unchanged or decreased across tumor types, with levels always remaining below those observed in heart mitochondria where OXPHOS demand is presumably highest.

**Fig. 1.**
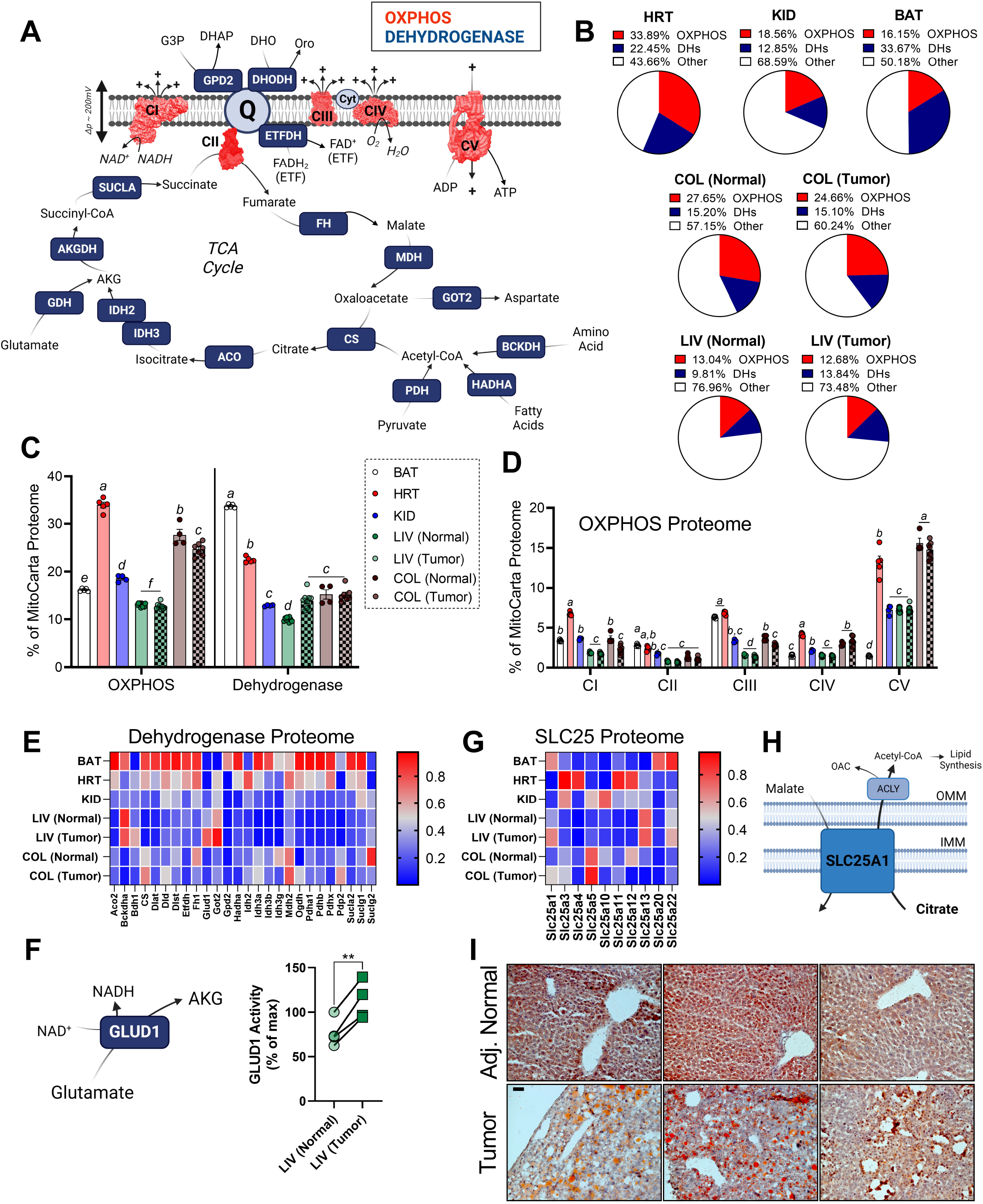
Pan-tissue analysis of the OXPHOS proteome in both normal and malignant mouse tissues. (**A**) Schematic of tissues analyzed, as well as the biochemical pathways incorporated in the analysis. (**B**) Pie charts depicting the percentage of the mitochondrial proteome devoted to protein components of either the OXPHOS complexes, or the TCA cycle and associated dehydrogenases. (**C**) Sample specific quantification of the summary data depicted in panel B. (**D**) Percentage of the mitochondrial proteome devoted to each OXPHOS complex. (**E**) Heatmap depicting protein abundance of the TCA cycle proteome. Data expressed as a percentage of max for each quantified protein. (**F**) GLUD1 reaction mechanism and activity in isolated mitochondria. Data displayed as a percentage of max GLUD1 activity in matched normal liver. (**G**) Heatmap depicting protein abundance of the SLC25 family proteome. Data expressed as a percentage of max for each quantified protein. (**H**) SLC25A1 reaction – electroneutral exchange of malate for citrate. Once in the cytosol, ATP citrate lyase (ACLY) converts citrate to oxalacetate (OAC) and acetyl-CoA to support lipid biosynthesis. (**I**) Images (20x) of oil red O staining for neutral lipid and lipid droplets in flash frozen sections of liver tumors and matched adjacent normal liver. Scale bar is 50μm. Data are mean ± SEM, N=4-10/group, *P<0.05. BAT = brown adipose tissue; HRT = heart; KID = kidney; LIV (Normal) = adjacent normal appearing liver; LIV (Tumor) = liver tumors induced by one-time exposure to diethylnitrosamine in 14-day old mice; COL (Normal) = normal colon; COL (Tumor) = tumors induced by AOM/DSS exposure; IMM = Inner mitochondrial membrane; OMM = Outer mitochondrial membrane. (**C, D, E, G**) Analysis by Two-way ANOVA with Tukey’s multiple comparison’s test. (**F, H**) Analysis by t-test.

### Intrinsic elevations in glutamate metabolizing enzymes characterize mitochondria in mouse liver tumors

Like OXPHOS complex expression, similar pan-tissue heterogeneity was apparent in the matrix dehydrogenase network, with liver being the lowest and brown adipose tissue being the highest (**Fig. 1B-C**). With respect to dehydrogenase differences in normal vs cancer, results were cancer-specific, with colon tumors being unchanged and liver tumors being higher (**Fig. 1B-C**). Interestingly, when evaluating the individual expression of each DH enzyme in the panel, enzymes related to glutamate metabolism were uniquely upregulated in liver tumors (**Fig. 1E**). Specific enzymes included aspartate aminotransferase (GOT2) and glutamate dehydrogenase (GLUD1). As validation of the proteomics results, the activity of GLUD1 was elevated in liver tumor mitochondria (**Fig. 1F**). Importantly, both GOT2 and GLUD1 expression were higher in liver tumors compared to adjacent normal liver, as well as all other tissues. Given the uniquely high expression/function of GLUD1 in liver tumors, it is conceivable that targeting glutamate metabolism in HCC may provide a superior therapeutic window over strategies that attempt to inhibit mitochondrial metabolism at more ubiquitously expressed targets (e.g., complex I).

### Analysis of the SLC25 mitochondrial carrier family reveals common upregulations in the citrate exporter SLC25a1 in mouse tumors

Metabolite exchange across the inner mitochondrial membrane is mediated by a family of transmembrane carriers known as the SLC25 mitochondrial carrier family^34^. Percent SLC25 expression varied widely across tissue types (**Supplemental Table 1**), consistent with varied energetic and biosynthetic demands across tissues. Restricting analysis to shared SLC25 family members, relative to matched normal, both colon and liver tumors diverted a greater percentage of their mitochondrial proteomes to the malate/citrate exchanger SLC25a1 (**Fig. 1G**). SLC25a1 catalyzes the electroneutral export of citrate to the cytosol to fuel lipid biosynthesis (**Fig. 1H**). As has been reported in CRC^35^, high SLC25a1 expression suggests that enhanced citrate export may facilitate increased de novo lipogenesis in liver cancer. Consistent with this, relative to matched normal, neutral lipid staining with oil red O revealed numerous large lipid deposits in liver tumors (**Fig. 1I**).

### Quantitative determination of respiratory capacity per unit of mitochondrial protein

Having established that mitochondrial proteome composition varies widely across both normal and malignant mouse tissues, we next wanted to investigate changes in respiratory flux. To do this, we made use of a technique that integrates in situ functional readouts of mitochondrial flux with quantitative determination of mitochondrial content using mass-spectrometry, effectively allowing for measurements of both respiratory capacity and OXPHOS kinetics to be interpreted both on a per cell and per mitochondrion basis^25,36–38^. Experiments were performed in permeabilized colorectal cancer (CRC) cells (e.g., CT26.WT) with results being compared to either permeabilized mouse colon or permeabilized heart myofibers (**Fig. 2A**). To assess respiratory capacity, plasma membrane permeabilization was first induced via exposure to cholesterol-specific detergents (e.g., saponin or digitonin). Carbon substrates were then added to maximize both NADH and FADH_2_ redox poise and maximal respiration was assessed via titration of the respiratory uncoupler FCCP. Across all cell types, exogenous cytochrome C did not impact respiration, indicating sustained integrity of the outer mitochondrial membrane (**Fig. 2B**; compare *J*O_2_ with ‘Pyr/M’ vs ‘Cyt C’). Relative to both normal colon and CRC cells, respiratory capacity was > 4-fold higher in permeabilized heart myofibers (**Fig. 2B-C**). Consistent with higher respiratory capacity in the heart, mitochondrial content, quantified by the ratio of mitochondrial protein/total protein abundance (i.e., mitochondrial enrichment factor; MEF), was also highest in heart myofibers (**Fig. 2D**). Mitochondrial content was equivalent in permeabilized normal colon strips relative to CRC cells (**Fig. 2D**). Armed with paired quantification of both respiratory kinetics and mitochondrial content, respiration data were normalized to flux per mitochondrion by scaling respiration results to each sample’s MEF. Upon normalization to MEF, relative to both normal colon and CRC cells, respiratory capacity remained ∽2-fold higher in heart myofibers (**Fig. 3A-B**). Higher respiration per unit of mitochondrial protein in the heart is consistent with the heart diverting a greater percentage of the mitochondrial proteome to the components of both OXPHOS and the associated dehydrogenase network (**Fig. 1B**). Relative to normal colon, respiratory capacity normalized to mitochondrial protein was unchanged in CRC cells (**Fig. 3A-B**). Unaltered respiratory capacity in colon cancer cells agrees with prior work demonstrating no change in respiratory capacity in liver tumors^36^. Thus, integration of in situ bioenergetic flux with proteomics-based quantification of mitochondrial content indicates that intrinsic respiratory capacity is largely unaltered in mouse tumors.

**Fig. 2.**
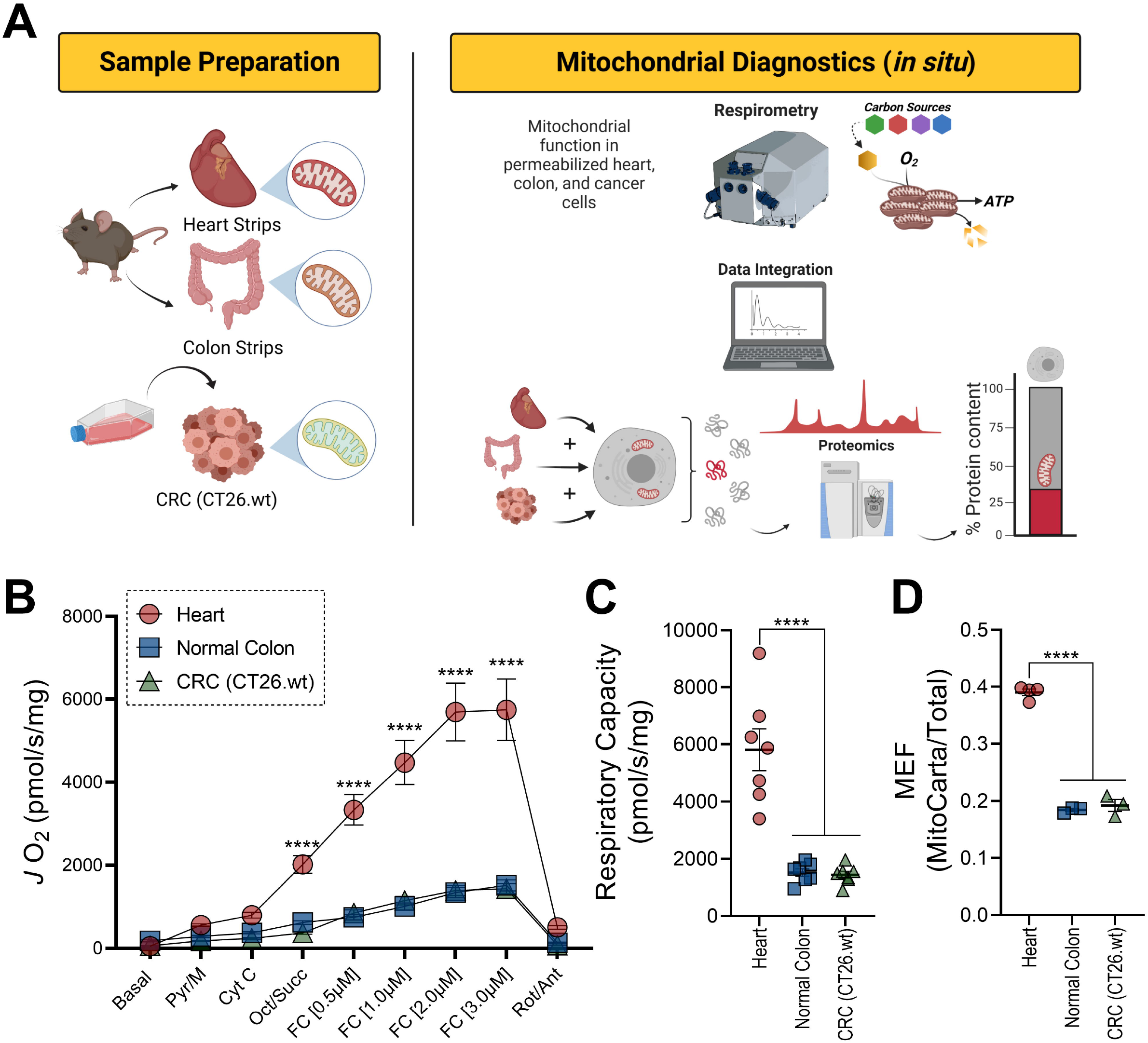
In situ quantification of respiratory capacity and mitochondrial content in mouse tissues. (**A**) Permeabilized tissue strips and cells were subjected to mitochondrial characterization using high resolution respirometry and proteomics analysis. (**B**) Maximal uncoupled respiration in permeabilized cells and tissues. Data normalized to total protein. (**C**) Quantified maximal respiratory capacity. (**D**). Ratio of mitochondrial protein to total protein abundance across samples. This ratio is referred to as Mitochondrial Enrichment Factor (MEF). Data are mean ± SEM, N=7/group (**B-C**), N=3-4/group (**D**), *P<0.05. Pyr = pyruvate; M = malate; Cyt C = cytochrome C; Oct = octanoyl-carnitine; Succ = succinate; FC = FCCP; Rot = rotenone; Ant = antimycin. (**B**) Analysis by Two-way ANOVA with Tukey’s multiple comparison’s test. (**C, D**) Analysis by One-way ANOVA with Tukey’s multiple comparison’s test.

**Fig. 3.**
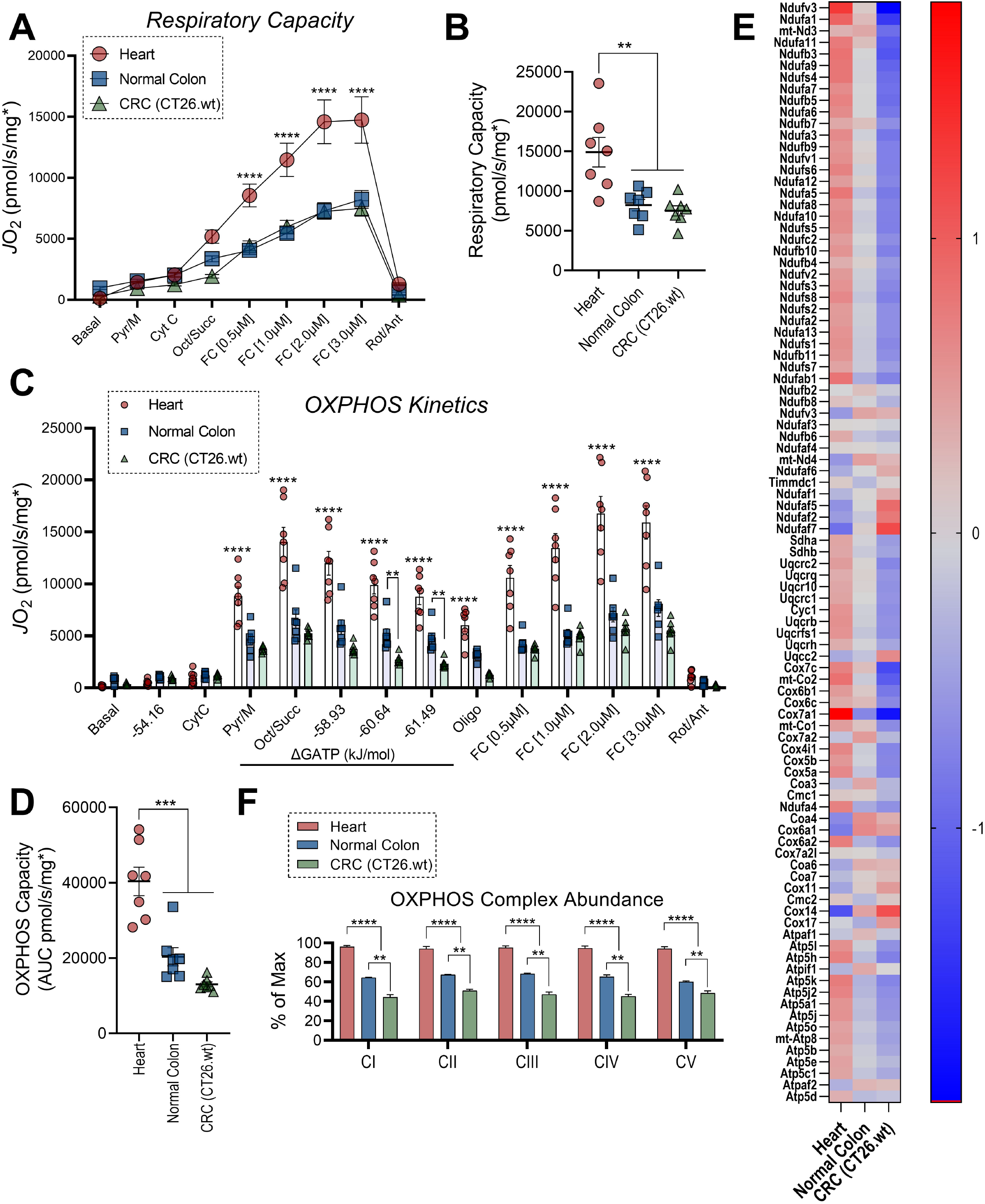
Lower intrinsic OXPHOS function in mouse colorectal cancer cells. (**A**) Maximal uncoupled respiration in permeabilized cells and tissues. Data normalized to total protein and then further normalized to mitochondrial protein using group specific MEF. (**B**) Quantified maximal respiratory capacity. (**C**) Assessment of OXPHOS kinetics using the CK clamp technique. (**D**) Quantified OXPHOS capacity, expressed as the area under curve during ΔG_ATP_ titration. (**E**) Heatmap depicting normalized protein abundance for the individual subunits that make up the OXPHOS complexes. (**F**) Summed abundance of individual complexes. Data displayed as a percentage of maximum abundance measured across all samples. Data are mean ± SEM, N=7/group (**A-D**), N=3-4/group (**E-F**), *P<0.05. Pyr = pyruvate; M = malate; Cyt C = cytochrome c; Oct = octanoyl-carnitine; Succ = succinate; FC = FCCP; Oligo = oligomycin; Rot = rotenone; Ant = antimycin. (**A, C, F**) Analysis by Two-way ANOVA with Tukey’s multiple comparison’s test. (**B, D**) Analysis by One-way ANOVA with Tukey’s multiple comparison’s test.

### Intrinsic deficiencies in OXPHOS characterize the mitochondrial network across distinct tumor types

The process of oxidative ATP synthesis functions as a ‘primed’ engine, whereby OXPHOS flux continuously adjusts to meet the prevailing demand for ATP regeneration (i.e., the energetic set point of cellular ATP free energy – i.e., ΔG_ATP_). To explore intrinsic OXPHOS functionality in cancer, we assessed respiratory control across a physiological span of ΔG_ATP_ (mimicking physiological transitions from high to low rates of ATP utilization). Titrations in the steady-state ΔG_ATP_ were made possible via a modified version of the creatine kinase (CK) energetic clamp^39–41^. This technique leverages the enzymatic activity of CK, which couples the interconversion of ATP and ADP to that of phosphocreatine (PCr) and free creatine (Cr) to titrate the extra-mitochondrial ATP/ADP ratio. Following activation of the OXPHOS system, carbon substrates were sequentially added to assess substrate/complex dependent (i.e., CI vs CI+CII) maximal OXPHOS flux, followed by titration of extramitochondrial ATP/ADP, inhibition of OXPHOS flux with oligomycin, and FCCP titration. Respiration coupled to OXPHOS, as well as that stimulated by the uncoupler FCCP, were several-fold higher in heart myofibers relative to normal colon and CRC cells (**Fig. 3C-D**). Relative to normal colon, despite no differences in respiration at more negative ATP free energies, increasing the extramitochondrial ATP/ADP revealed lower respiration in CRC cells, suggestive of reduced OXPHOS functionality in CRC (**Fig. 3C**). Consistent with high OXPHOS in heart and lower OXPHOS flux in CRC, percent expression of all five OXPHOS complexes was highest in the heart and lowest in CRC cells (**Fig. 3E-F**).

Turning our attention to another tumor type, in situ mitochondrial phenotyping was carried out in digitonin-permeabilized leukemia cells originally isolated from C57BL/6J mice with acute myeloid leukemia (AML)^42^. Results were compared to bone marrow mononuclear cells (BMMC) isolated from long bones of C57BL/6J mice. Normalized to total protein, respiratory capacity was similar between BMMC and AML (**Fig. 4A**). Despite similar respiratory capacities, total mitochondrial protein abundance was higher in AML, indicative of higher mitochondrial content (**Fig. 4B-C**). Like liver and colon tumors, the percentage of the mitochondrial proteome dedicated to OXPHOS, and the dehydrogenase network was largely similar in AML and BMMC (**Fig. 4D**). Breaking down OXPHOS abundance based on each OXPHOS complex, CI was slightly elevated in AML, with both CIII and CIV being lower in AML (**Fig. 4E-F**). Relative to BMMC, despite no differences in OXPHOS respiration when supported by NADH-linked pyruvate/malate, saturation of the carbon substrate pool with fatty acid and succinate revealed lower OXPHOS flux in AML (**Fig. 4G**). Thus, although mitochondrial content is elevated in AML, intrinsic deficiencies in the distal OXPHOS complexes (e.g., CIII and CIV) restrict OXPHOS flux in AML. Lower intrinsic OXPHOS function in mouse AML is similar to that previously reported in human AML^43^, as well as that observed herein for both liver and colon tumors. In conclusion, comprehensive bioenergetic phenotyping of cancerous mitochondria indicates that for a fixed amount of mitochondrial protein, OXPHOS function is generally downregulated across tumor types.

**Fig. 4.**
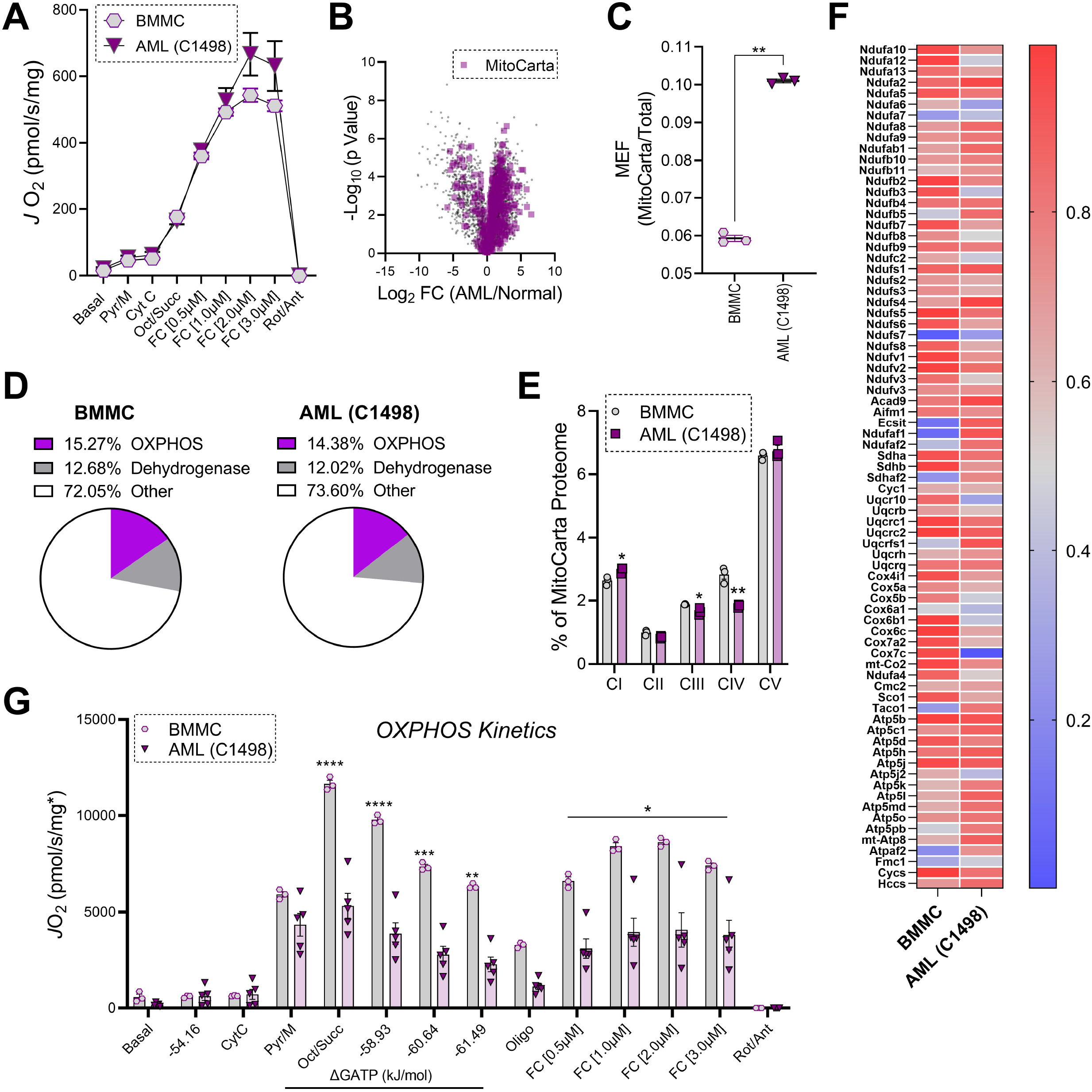
Lower intrinsic OXPHOS function in mouse acute myeloid leukemia cells. (**A**) Maximal uncoupled respiration in permeabilized mouse BMMC and AML cells. Data normalized to protein. (**B**) Volcano pot depicting proteomic changes between normal and leukemic blood cells with mitochondrial proteins shown in purple. (**C**) Ratio of mitochondrial protein to total protein abundance across samples (MEF). (**D**) Pie charts depicting the percentage of the mitochondrial proteome devoted to protein components of either the OXPHOS complexes, or the TCA cycle and associated dehydrogenases. (**E**) Percentage of the mitochondrial proteome devoted to each OXPHOS complex. (**F**) Heatmap depicting normalized protein abundance for the individual subunits that make up the OXPHOS complexes. (**G**) Assessment of OXPHOS kinetics using the CK clamp technique. Data normalized to mitochondrial enrichment factor. Data are mean ± SEM, N=3/group (**B-F**), N=3-5/group (**A, G**), *P<0.05. BMMC = bone marrow mononuclear cells; AML = acute myeloid leukemia; Pyr = pyruvate; M = malate; Cyt C = cytochrome c; Oct = octanoyl-carnitine; Succ = succinate; FC = FCCP; Oligo = oligomycin; Rot = rotenone; Ant = antimycin. (**A, C, E-G**) Analysis by t-test.

## DISCUSSION

Although all mitochondria make ATP, emerging evidence indicates that the demand for OXPHOS, as well as the various other mitochondrial outputs (ROS, NADPH, metabolites), varies widely across organ systems^11,27,39^. Simply put, mitochondria across the body’s > 200 cell types are highly heterogeneous. At present, it appears that tissue-specific differences in mitochondrial function (i.e., mitochondrial specialization) are operationally defined through differences in mitochondrial protein expression, as well as an ever-growing list of post-translational modifications^44–47^. Regardless of the mechanism(s), the biological reward of mitochondrial specialization is the alignment of bioenergetic function with organ physiology (i.e., establishment of bioenergetic fidelity). In the case of cancer, bioenergetic fidelity becomes misaligned from the host tissue resulting in neoplastic growth. While this misalignment is in part associated with increased glucose uptake/glycolytic flux, multiple lines of evidence link various aspects of cancer biology (e.g., tumorigenesis, chemoresistance) to altered mitochondrial quantity and quality^36,38,43,48–54^. Collectively, these studies lend strong support to the notion that current endeavors into metabolism-tailored pharmacotherapies should consider targeting the mitochondria. While these studies have ignited interest in mitochondrial-targeted chemotherapeutics, clinical success for a given ‘mito-therapeutic’ will undoubtedly hinge upon its ability to specifically target only cancerous mitochondria.

With respect to targeting mitochondrial oxidative metabolism in cancer, a handful of drugs have entered clinical trials. Drugs that have been trialed or are currently being trialed include CPI-613, which targets the lipoate-dependent dehydrogenase complexes for pyruvate, alphaketoglutarate, and branched-chain ketoacids; glutaminase inhibitors; as well as inhibitors of respiratory complex I (e.g., metformin and IACS-010759)^23,54–56^. Although complex I has been demonstrated to be essential for tumor growth^10,13^, when normalized to a fixed amount of mitochondrial protein, expression of complex I was unchanged in liver tumors and decreased in colon tumors. Although slight elevations in normalized complex I expression were apparent in AML cells relative to BMMC, complex I levels in AML cells were still 2-fold lower than that of heart mitochondria. For example, heart mitochondria divert nearly 7% of their mitochondrial proteome to protein subunits of complex I, suggesting that maintaining energy homeostasis in high-OXPHOS-demand organs like heart requires enhanced complex I function. Interestingly, mutations in mtDNA-encoded complex I genes are frequently observed in human tumors^58,59^. Given that impaired complex I function has been observed in several cancer types^6,36,60–63^, including colon and liver tumors of mice^36^, such findings raise the intriguing hypothesis that ‘restoring’, rather than ‘inhibiting’ complex I function may represent an untapped anti-cancer strategy.

Mitochondrial content varies widely by cell type and is highly influenced by cellular stress. For example, lesions to OXPHOS can induce compensatory upregulations in mitochondrial content via retrograde signaling^64,65^, as exemplified in renal oncocytomas driven by genetic lesions in complex I^61,66,67^. Thus, experiments designed to investigate the role of the mitochondria in cancer would seemingly benefit from techniques capable of normalizing both subcellular proteomic and bioenergetic flux data to mitochondrial content. To this end, we recently developed a label-free proteomics method capable of quantitating percent mitochondrial enrichment on a per-sample basis by comparing mitochondrial protein intensity (e.g., the sum of all MitoCarta+ proteins) to total protein intensity^25^. This generates a ‘mitochondrial enrichment score’ that normalizes proteomics data and bioenergetic flux readouts to defined amounts of mitochondrial protein^25^, enabling direct comparisons of mitochondrial networks across distinct cell types (i.e., tumor types and matched normal tissue). Relative to a fixed amount of mitochondrial protein, protein components of the OXPHOS complexes, as well as OXPHOS flux were consistently observed to be downregulated across tumor types. Although reliance on mouse cancer cell lines and early-stage in vivo tumor models limits the translation of our findings to human tumors, lower intrinsic OXPHOS function in tumors provides potential insight into the systemic toxicity observed with indiscriminate OXPHOS inhibitors in human cancer trials^24^.

Effective cancer therapeutics are drugs that target biochemical pathways unique to cancer cells. However, cancer cells use a highly similar repertoire of proteins and signaling pathways as normal cells^68^, making identification of cancer-specific features a significant barrier. Recent efforts have begun to overcome this barrier by exploiting normal versus cancer cell biochemical differences that originate within the mitochondria^36,43,54,69^. Although mitochondria are present in all cells (red blood cells excluded), mitochondria from different cell types are remarkably unique in both composition and function, especially in cancer cells. For example, independent of OXPHOS demand/reliance, considerable heterogeneity was observed herein across tumor types with respect to the stoichiometry of the subunits comprising the OXPHOS complexes, individual dehydrogenase enzymes (e.g., glutamate metabolism in HCC), as well as SLC25 mitochondrial transporters (e.g., SLC25a1). Although the potential utility of leveraging these cancer-specific bioenergetic features to combat the disease remains to be determined, the present results raise the exciting potential for cancer cells’ unique bioenergetic signatures to be leveraged to design chemotherapeutic approaches that specifically target only cancer cell mitochondria. The first step to realizing the goal of mitochondrial precision medicine is the identification of tumor-type specific mitochondrial phenotypes.

## METHODS

### Materials

All chemicals for cell culture and animal treatments, as well as mitochondrial and proteomic analysis, were purchased from either Millipore Sigma or Thermo Fisher Scientific. Fetal bovine serum (FBS) was purchased from Peak Serum (Wellington, CO).

### Cell Culture

Murine colorectal carcinoma cells (CT26.WT) were purchased from ATCC (#CRL-2638). CT26.WT cells were cultured in RPMI-1640 medium supplemented with 10% FBS and 1% penicillin/streptomycin. Prior to cell harvest, cells were seeded in T75 culture flasks and allowed to reach confluency. Once confluent, cells were harvested with Trypsin-EDTA (0.25%). Murine acute myeloid leukemia cells (C1498) were originally purchased from ATCC (#TIB-49). C1498 cells used in this study were provided by Dr. Tan (UVA). C1498 cells were cultured in RPMI-1640 medium supplemented with 10% FBS and 1% penicillin/streptomycin. Cells were monitored daily and harvested when the average cell density reached 1.0 × 10^6^ cells/mL.

### Animal Treatment

All procedures on experimental animals were approved by the East Carolina University Institutional Animal Care and Use Committee. Mice were housed under controlled temperature (22.7°C) and light (12h light/12h dark) conditions with free access to food and water. Proteomics data in Figure 1 pertaining to isolated mitochondria from brown adipose tissue (BAT), heart (HRT), and kidney (KID) were analyzed from a previously published dataset^25^. In the prior study^25^, tissues were harvested from C3H/HeJ mice (n=10; The Jackson Laboratory, stock #000659) after a 12-hour fast. Proteomics data in Figure 1 pertaining to isolated mitochondria from liver tumors (LIV Tumor) and matched normal liver tissue (LIV Normal) were analyzed from a previously published dataset^36^. Induction of hepatocellular carcinoma was achieved using a single injection of diethylnitrosamine (DEN), as previously described^70^. DEN (25mg/kg) or saline vehicle (0.9% NaCl) was injected intraperitoneally at 14 days postnatal. As mice were not weaned, all littermates received the same treatment. Both DEN and saline mice were sacrificed after tumor development at ∽27 weeks of age. At the time of sacrifice, 12h-fasted mice were anesthetized with isofluorane and exsanguinated prior to tissue removal. Induction of colorectal cancer was performed in C57BL/6J mice (Jackson Laboratory, stock #000664). To induce colorectal tumors, 8-12-week-old male and female mice received a single I.P. injection of azoxymethane (AOM; 8mg/kg) dissolved in PBS. One week following AOM injection, mice were exposed to dextran sodium sulfate (DSS; 3%) in the drinking water for a period of 5 days. Mice were allowed to recover with normal water for a period of 14 days. This cycle of DSS exposure (5 days on, 14 days off) was repeated for a total of three times. The AOM/DSS exposure was modeled after prior publications^63,71^. Mice were sacrificed 90-95 days post AOM injection and colorectal tumors were harvested for experiments. For normal colon, sections of colon were collected from age and sex matched C57BL/6J mice. Given that CT26.WT cells were isolated from BALB-cJ mice, experiments in Figures 2-3 pertaining to in situ mitochondrial phenotyping were performed in tissues (heart and colon) harvested from BALB-cJ mice (Jackson Laboratory, stock #000651). Given that C1498 cells were isolated from C57BL/6J mice, experiments in Figure 4 pertaining to in situ mitochondrial phenotyping were performed in bone marrow mononuclear cells (BMMC) harvested from C57BL/6J mice. To isolate BMMC, male and female C57BL/6J mice were anesthetized with isofluorane and exsanguinated prior to removal of the long bones. Bone marrow was collected via aspiration with RPMI-1640 using a 26G needle. Cells were treated with ACK lysis buffer to remove red blood cells. ACK-treated cells were then passed through a 70μm cell strainer, pelleted at 500 x g for 10 minutes at room temperature. Cell pellets were then suspended in the indicated experimental buffer and used for either bioenergetic or proteomic analysis.

### Isolation of mitochondria from mouse tissues

Mitochondria were isolated from tissues using differential centrifugation as described previously, with some modifications^72^. The following buffers were utilized: Buffer A - MOPS (50mM), KCl (100mM), EGTA (1mM), MgSO_4_ (5mM), pH=7.1; Buffer B - Buffer A, supplemented with bovine serum albumin (BSA; 2g/L). After removal, all tissues were immediately placed in ice-cold Buffer B, minced, then homogenized via a drill-driven Teflon pestle and borosilicate glass vessel. Homogenates were centrifuged at 800 x g for 10min at 4°C. The supernatant was filtered through gauze and centrifuged at 10,000 x g for 10min at 4°C. The pellets were washed in 1.4mL of Buffer A, transferred to microcentrifuge tubes, and again centrifuged at 10,000 x g for 10min at 4°C. Final mitochondrial pellets were resuspended in 100-200mL of Buffer A. Protein content was determined via the Pierce BCA protein assay.

### Proteomics sample preparation

Samples used for proteomics analysis included the following tissues: **A**) isolated mitochondria from BAT, HRT, KID, LIV (Tumor), LIV (Normal), COL (Normal) – Figure 1; **B**) Intact tissues from colon tumors (COL-Tumor) and normal colon (COL-Normal) – Figure 1; **C**) Saponin/digitonin permeabilized tissues or cells (Heart, Normal Colon, CRC (CT26.wt)) - Figures 2-3; **D**) Intact cell pellets (BMMC, AML (C1498)) – Figure 4. All samples were lysed and digested as previously described^73^. Briefly, samples were lysed in urea lysis buffer (8M urea in 40mM Tris, 30mM NaCl, 1mM CaCl_2_, 1 cOmplete ULTRA mini EDTA-free protease inhibitor tablet; pH=8.0), subjected to two freeze-thaw cycles and sonication for 5s at an amplitude of 30 using a probe sonicator (Q Sonica #CL-188). Equal amounts of protein (100μg) were then reduced with 5mM dithiothreitol (30min incubation at 32°C), alkylated with 15mM iodoacetamide (30min incubation in the dark at room temperature), and then unreacted iodoacetamide was quenched with an additional 10mM dithiothreitol. Digestion was performed in two steps: first using Lys C (ThermoFisher, Cat#90307; 1:100 w:w enzyme:protein; 4h incubation at 32°C), followed by dilution of the samples to 1.5M urea using 40mM Tris (pH=8.0), 30mM NaCl, 1mM CaCl_2_ for overnight digestion via trypsin at 32°C (Promega, Cat# V5113; 1:50 w:w enzyme:protein). Following digestion, samples were acidified to 0.5% TFA, centrifuged at 10,000 x g for 10min to pellet undigested material, and the supernatant was collected for desalting of soluble peptides using 50mg tC18 SEP-PAK solid phase extraction columns (Waters; Cat# WAT054955) as previously described^74^. The resulting eluate was frozen on dry ice and subjected to speedvac concentration.

### TMT labeling

Quantification was done using Tandem Mass Tags (TMT) plex for proteome characterization of permeabilized tissues (Figures 2-3). TMT labeling was performed as previously described^43^. Peptides were suspended in 100□μL of 200□mM triethylammonium bicarbonate (TEAB), mixed with a unique 10-plex Tandem Mass Tag (TMT) reagent (0.8□mg re-suspended in 50 μL100% acetonitrile), and shaken for 4□hr at room temperature (Thermo Fisher). The TMT reagent labeling strategy was as follows: 126-Colon, 127N-Heart, 127C-Heart, 128N-Colon, 128C-Colon, 129N-Heart, 129C-Heart, 130N-CT26.WT, 130C-CT26.WT, 131-CT26.WT. Following quenching with 0.8□μL 50% hydroxylamine, samples were frozen, and subjected to speedvac concentration. Samples were suspended in ∽1□mL of 0.5% TFA and again subjected to solid phase extraction, but with a 100□mg tC18 SEP-PAK SPE column (Waters). The multiplexed peptide sample was subjected to high pH-reversed-phase fractionation according to the manufacturer’s instructions (Thermo Fisher). In this protocol, peptides (100□μg) are loaded onto a pH-resistant resin and then desalted with water washing combined with low-speed centrifugation. A step-gradient of increasing acetonitrile concentration in a high-pH elution solution is then applied to columns to elute bound peptides into 8 fractions. Following elution, fractions were frozen and subjected to speedvac concentration.

### nLC-MS/MS

Final peptides were resuspended in 0.1% formic acid for peptide quantification (ThermoFisher Cat# 23275) and dilution to a final concentration of 0.25μg/μL. nanoLC-MS/MS analysis was performed using an UltiMate 3000 RSLCnano system (ThermoFisher) coupled to a Q Exactive Plus Hybrid Quadrupole-Orbitrap mass spectrometer (ThermoFisher) via a nanoelectrospray ionization source as previously described ^73^. MS1 was performed at a resolution of 70,000, with an AGC target of 3×10^6^ ions and a maximum injection time (IT) of 100ms. Data-dependent acquisition (DDA) was used to collect MS2 spectra of the top 15 most abundant precursor ions with a charge >1 per MS1 scan, with dynamic exclusion enabled for 20s. The isolation window for precursor ions was 1.5m/z, and normalized collision energy was 27. MS2 scans were performed at 17,500 resolution, maximum IT of 50ms, and AGC target of 1×10^5^ ions.

As previously described^43^, for TMT-labeled samples, peptide fractions were suspended in 0.1% formic acid at a concentration of 0.25□μg/μL, following peptide quantification (ThermoFisher). All samples were subjected to nanoLC-MS/MS analysis using an UltiMate 3000 RSLCnano system (Thermo Fisher) coupled to a Q Exactive PlusHybrid Quadrupole-Orbitrap mass spectrometer (Thermo Fisher) via nanoelectrospray ionization source. For each injection of 4□μL (1□μg), the sample was first trapped on an Acclaim PepMap 100 20□mm × 0.075□mm trapping column (Thermo Fisher) 5□μl/min at 98/2□v/v water/acetonitrile with 0.1% formic acid, after which the analytical separation was performed over a 90-min gradient (flow rate of 300 nanoliters/min) of 3 to 30% acetonitrile using a 2□μm EASY-Spray PepMap RSLC C18 75□μm × 250□mm column (Thermo Fisher) with a column temperature of 55□°C. MS1 was performed at 70,000 resolution, with an AGC target of 1 × 10^6^□ions and a maximum IT of 60□ms. MS2 spectra were collected by data-dependent acquisition (DDA) of the top 20 most abundant precursor ions with a charge greater than 1 per MS1 scan, with dynamic exclusion enabled for 30s. Precursor ions were filtered with a 1.0□m/z isolation window and fragmented with a normalized collision energy of 30. MS2 scans were performed at 17,500 resolution, AGC target of 1 × 10^5^□ions, and a maximum IT of 60□ms.

### Data analysis for label-free quantitative proteomics

Proteome Discoverer 2.2 (PDv2.2) was used for raw data analysis. Default search parameters included oxidation as a variable modification and carbamidomethyl (57.021 Da on C) as a fixed modification. Data were searched against both the Uniprot Mus musculus reference proteome and mouse Mito Carta 3.0 database^75^. As previously described^73^, PSMs were filtered to a 1% FDR and grouping to unique peptides was also maintained at a 1% FDR at the peptide level. Strict parsimony was used to group peptides to proteins, and proteins were again filtered to 1% FDR. MS1 precursor intensity was used for peptide quantification, and low abundance resampling was used for imputation. As previously described^73^, high-confidence master proteins were used to determine mitochondrial enrichment factor (MEF) by quantifying the ratio of mitochondrial protein abundance (identified using the MitoCarta 3.0 database) to total protein abundance. Sample-specific MEF was then used to first normalize all proteomics data to total mitochondrial protein abundance. Further normalization was performed by converting individual abundances of each quantified mitochondrial protein to percent contribution to total mitochondrial abundance. For quantification of the OXPHOS complexes, we summed the percent contribution of all subunits quantified that match to MitoCarta 3.0 annotated subunits of CI, CII, CIII, CIV, and CV.

### Data analysis for TMT quantitative proteomics

Proteome Discoverer 2.2 (PDv2.2) was used for raw data analysis, with default search parameters including oxidation (15.995□Da on M) as a variable modification and carbamidomethyl (57.021□Da on C) and TMT6plex (229.163□Da on peptide N-term and K) as fixed modifications, and 2 missed cleavages (full trypsin specificity). Data were searched against both the Uniprot Mus musculus reference proteome and mouse Mito Carta 3.0 database^75^. PSMs were filtered to a 1% FDR. PSMs were grouped to unique peptides while maintaining a 1% FDR at the peptide level. Peptides were grouped to proteins using the rules of strict parsimony and proteins were filtered to 1% FDR using the Protein FDR Validator node of PD2.2. MS2 reporter ion intensities for all PSMs having co-isolation interference below 0.5 (50% of the ion current in the isolation window) and an average S/N > 10 for reporter ions were summed together at the peptide and protein level. Imputation was performed via low abundance resampling. The protein group tab in the PDv2.2 results was exported as tab delimited.txt. files, and analyzed based on a previously described workflow^43^. M2 reporter (TMT) intensities were converted to log_2_ space and the average value from the ten samples per kit was subtracted from each sample specific measurement to normalize the relative measurements to the mean of each master protein. For protein-level quantification, only Master Proteins—or the most statistically significant protein representing a group of parsimonious proteins containing common peptides identified at 1% FDR—were used for quantitative comparison.

### In situ analysis of mitochondrial respiration

High-resolution O_2_ consumption measurements were conducted using the Oroboros Oxygraph-2K (Oroboros Instruments, Innsbruck, Austria). All experiments were carried out at 37°C in a 1-mL reaction volume using Respiration Buffer (potassium-MES; 105mM; pH 7.2, KCl; 30mM), KH_2_PO_4_; 10mM, MgCl_2_; 5mM, EGTA; 1mM, creatine monohydrate; 5mM, and BSA; 2.5 g/L). The following procedures relate to permeabilized tissue and cell experiments detailed in Figures 2-3. Cardiac myofibers and sections of colon mucosa were subjected to a combination of mechanical (fine forceps under a dissecting scope) and chemical permeabilization (saponin; 30μg/mL) in Respiration Buffer. Permeabilized tissues were then washed in Respiration Buffer and placed inside the O2K chambers for analysis. For CT26.WT cells, permeabilization was performed in cells suspended at 1 × 10^6^ cells/mL in Respiration Buffer supplemented with 0.015μg/mL digitonin. Cells were permeabilized for 30 minutes on a rocker at 4°C. Following permeabilization, cells were pelleted at 500 x g for 10 minutes at 4°C. Cell pellets were suspended in Respiration Buffer and placed inside the O2K chambers for analysis. For respiration experiments, Respiration Buffer was supplemented with 10μM blebbistatin to prevent contraction of the cardiac myofibers. Following analysis, tissues and cells were collected and lysed in Urea lysis buffer to allow for normalization to total protein. Total protein determined via the Pierce BCA protein assay. The following procedures relate to permeabilized cell experiments detailed in Figure 4. BMMC and C1498 cells were suspended in Respiration Buffer at a cell concentration of 1-3 × 10^6^ cells/mL. Cell permeabilization was performed inside the O2K chambers using 0.015μg/mL digitonin. Following respiration analysis, cells were collected, pelleted at 500 x g for 10 minutes at 4°C, and lysed in 1x RIPA lysis buffer to allow for normalization to total protein. Total protein was determined via the Pierce BCA protein assay.

For experiments designed to assess respiratory capacity, after recording basal respiration, samples were energized with various carbon substrates (pyruvate, malate, succinate, octanoyl-l-carnitine; Pyr, M, Succ, Oct; 5 mM, 2 mM, 5 mM, 0.2 mM) and flux was stimulated with FCCP titration (0.5-3.0μM). Cytochrome C (10μM) was added to check the integrity of the outer mitochondrial membrane. Note, the absence of an increase in respiration, relative to the pre cytochrome C rate, was used as a quality control assessment for outer membrane integrity. A combination of rotenone (500nM) and antimycin (500nM) was added at the end of the protocol to control for non-mitochondrial oxygen consumption. OXPHOS kinetics was assessed across a physiological ATP free energy demand using the creatine kinase (CK) clamp as previously described^25,26^. For complete details regarding the calculation of ΔG_ATP_ at each titration point see Fisher-Wellman et al^26^. OXPHOS kinetics experiments also included the addition of the ATP synthase inhibitor oligomycin (20nM) to determine ATP-linked respiration. Data were normalized to total protein and then corrected for the mitochondrial enrichment factor (MEF) calculated for each group (see proteomics methodology above regarding MEF). All additions were made directly to the O2K chamber during the period of each assay. Typical assay length was 20-40 minutes.

### GLUD1 activity assay

Enzyme activity of GLUD1 (glutamate dehydrogenase) was determined via the autofluorescence of NADH (Ex:Em 340:450), as described previously^26^. Assays were performed in Respiration Buffer using freeze-fractured isolated mitochondria from liver tumors or matched normal liver tissue. Following the addition of glutamate, fluorescence was measured every minute for 20 minutes in the plate reader.

### Histological analysis of liver tissues

Flash-frozen sections of liver tumors and matched normal liver were embedded in Optimal Cutting Temperature compound (O.C.T) and then flash-frozen using liquid N2-chilled isopentane. Blocks were sections at 12μm thickness and used to detect neutral lipids and lipid droplets via Oil red O staining. Slides were incubated in 4% paraformaldehyde for 5 minutes, rinsed in distilled water, subjected to 2 × 3-minute incubations in 100% ethylene glycol, then exposed to oil-Red-O for 20 minutes. Following oil-red-O exposure, slides were exposed to 85% ethylene glycol for 3 minutes, rinsed in distilled water, and exposed to Mayer’s hematoxylin for 5 minutes. Slides were developed under running tap water for 5 minutes and mounted with hot glycerin^76^. Images were acquired at 20X using a Zeiss Axio Observer microscope.

### Statistical analysis

For all proteomics data, the ‘protein’ tab in the PDv2.2 results was exported as a tab-delimited .txt. file and analyzed. Protein abundance was converted to the Log_2_ space. For pairwise comparisons, tissue mean, standard deviation, p-value (p; two-tailed Student’s t-test, assuming equal variance), and adjusted p-value (Benjamini Hochberg FDR correction) were calculated^77^. Mitochondrial functional assay results are expressed as the mean ± SEM (error bars). Data were normalized to mg total protein and then corrected for the group mean MEF, with the final values expressed as pmol/s/mg*, where the ‘*’ indicates protein values corrected to mitochondrial protein only using MEF^78^. Throughout the paper, differences between groups were assessed by t-test, as well as either one-way or two-way ANOVA, followed by Tukey’s test where appropriate using GraphPad Prism 8 software (Version 8.4.2). Other statistical tests used are described in the figure legends. Statistical significance in the figures is indicated as follows: \**p* < 0.05; \*\**p* < 0.01; \*\*\**p* < 0.001; \*\*\*\**p* < 0.0001. Figures were generated using GraphPad Prism 8 software (Version 8.4.2) or Biorender.

### Data availability

All raw data for proteomics experiments are available online using the following accession numbers “JPST000908”, “JPST002178”, “JPST002177”, “JPST001567”, “JPST002179” for jPOST Repository^79,80^. Analyzed proteomics data across all groups are presented in **Supplemental Table 1**. Source data for all figures are provided in **Supplemental Table 2**.

## Acknowledgments

This work was supported in part by DOD-W81XWH-19-1-0213 (K.H.F-W.), DK125812 (J.M.E., NIH/NIDDK), P01 CA171983 (T.P.L., M.K., M.C.C., NIH/NCI). Additional support for this project was provided in part by the Brody School of Medicine at ECU’s Mass Spectrometry Core which has received support from the Golden Leaf Foundation and federal COVID-19 relief funds appropriated to ECU in North Carolina SL 2020-2024.

## Conflicts of interest

The authors declare that the research was conducted in the absence of any commercial or financial relationships that could be construed as a potential conflict of interest.

## Supplementary Data

Supplemental **Table 1. Proteomics Data**. (**A**) OXPHOS complex and dehydrogenase database. (**B**) Analyzed results for Figure 1 – BAT, HRT, KID. (**C**) Analyzed results for Figure 1 – LIV (Tumor) and LIV (Normal). (**D**) Analyzed results for Figure 1 – COL (Normal), includes data from normal colon tissue, as well as isolated mitochondria from normal colon. (**E**) Analyzed results for Figure 1 – COL (Tumor). (**F**) Analyzed results for Figure 2 – Permeabilized tissues (Colon, CT26.WT, Heart). (**G**) Analyzed results for Figure 4 – AML (C1498), BMMC.

**Supplemental Table 2. Source Data**. Individual data points for all figures.

## Notes

### Competing Interest Statement

The authors have declared no competing interest.

